# Brain alignment in deep neural networks emerges early and independently of object classification

**DOI:** 10.64898/2026.08.18.745511

**Authors:** H. Steven Scholte, Niklas Müller, Julio Smidi, Iris I. A. Groen, Marcel A. J. van Gerven

## Abstract

Deep convolutional neural networks are leading models of biological vision, largely because of their strong brain alignment: their features predict neural responses better than earlier models. Yet they are believed to recognize objects differently, relying on texture where humans rely on shape and failing on perturbations humans handle effortlessly. What then does alignment reflect? Tracking three architectures densely during training, we find that alignment with human fMRI, EEG, and macaque electrophysiology is already largely present at initialization, when networks classify at chance, and reaches a plateau within one to five epochs; thereafter it changes only modestly while classification accuracy continues to climb to 75%. Network lesioning shows that a kernel’s contribution to alignment is essentially uncorrelated with its contribution to classification throughout training. Brain-network alignment therefore appears to reflect the structure of the visual environment both systems encode, rather than a shared solution for classification.

## Introduction

Deep convolutional neural networks (DCNNs) trained on object recognition are the leading computational and biologically plausible models of primate vision. Across the visual hierarchy, from V1 through inferotemporal cortex (IT), these networks predict neural responses better than non-deep learning models (Güçlü & Van Gerven, 2015; Kriegeskorte, 2015; Yamins & DiCarlo, 2016). This predictive success has been interpreted as evidence that biological and artificial vision systems converge on similar computational solutions: optimizing for object recognition yields representations like those found in brains (Yamins et al., 2014; Schrimpf et al., 2020).

Yet these same networks are believed to recognize objects in a fundamentally different way than humans do. Presented with images where shape and texture cues conflict, humans classify by shape while networks classify by texture (Geirhos et al., 2019). Networks rely on high frequency local patterns that humans largely ignore (Baker et al., 2018; Subramanian et al., 2023) and fail on image perturbations that humans handle effortlessly (Geirhos et al., 2018). In other words, the systems that best predict human brain activity do not behave like human observers.

If DCNNs solve object recognition differently than humans, what explains their alignment with human brain representations? The dominant explanation assumes a common goal: task optimization for core object recognition drives brain alignment (Yamins & DiCarlo, 2016). If this is correct, two predictions follow. First, brain alignment and object classification performance should improve together across training. Second, the same kernels should support both brain alignment and classification. Yet alignment has so far been assessed almost entirely for fully trained networks, as a function of architecture, training data, objective, and dimensionality (Doerig et al., 2023; Conwell et al., 2024), while how alignment itself evolves over training has received little attention. Following alignment across training is a direct test of these predictions: any alignment that is already present before a network can classify cannot be a product of task optimization.

We test these predictions by comparing a developing artificial system with developed biological vision: tracking DCNNs from initialization to convergence on ImageNet (Russakovsky et al., 2015) and measuring, at each checkpoint, how closely they resemble adult human and macaque neural recordings and behavior. We evaluate alignment against three large-scale neural datasets: 7T fMRI from human visual cortex (Allen et al., 2022), high density EEG (Gifford et al., 2022), and macaque ventral stream electrophysiology (Papale et al., 2025), together with behavioral measures of human similarity, generalization, and classification strategy (error consistency, out-of-distribution accuracy, and shape bias; Geirhos et al., 2018, 2019, 2020). This approach allows us to observe not just how well networks match biological vision, but how that correspondence develops, and when, during training, it peaks.

Our findings contradict both predictions. At initialization, when networks classify at chance, they already reach 62% of their maximum alignment, consistent with recent evidence that convolutional architectures are cortex-aligned de novo (Kazemian et al., 2025). We show that most of what remains is added within the first few epochs, while classification accuracy is still far from its final value, and the rest of training contributes only modestly. A kernel lesioning analysis reveals the mechanistic basis of this dissociation: the kernels that contribute to brain encoding are largely independent of those that support classification, with near-zero correlation throughout training. These results suggest that brain-network alignment does not reflect a shared solution to the classification objective but rather that both systems represent a shared natural image structure.

## Results

### Brain alignment rises and plateaus rapidly, within the first few training epochs

Throughout, we quantify brain alignment with linearized encoding models that predict neural responses from network activations (see Methods). For three neural datasets, we defined two regions or time points of interest, reflecting both lower and higher level visual processes: V1 and PPA (human fMRI), V1 and IT (macaque electrophysiology), and 120 and 200 ms (human EEG). For each of these neural targets (referred to as *early* and *late* targets), we built a separate encoding model using the extracted activations of a ResNet-50 at each saved checkpoint (see **Figure 1A**). **Figure 1B** and **1C** show the cross-validated encoding performances for each neural target alongside two behavioral measures (error consistency and out-of-distribution accuracy), across training. Of the three architectures we trained, ResNet-50 showed the highest average encoding performance across training (**Supplementary Figure 1**), motivating its use as our primary model; AlexNet and ConvNeXt-T are reported as generalization cases (**Supplementary Figures 2–5**).

**Figure 1.**
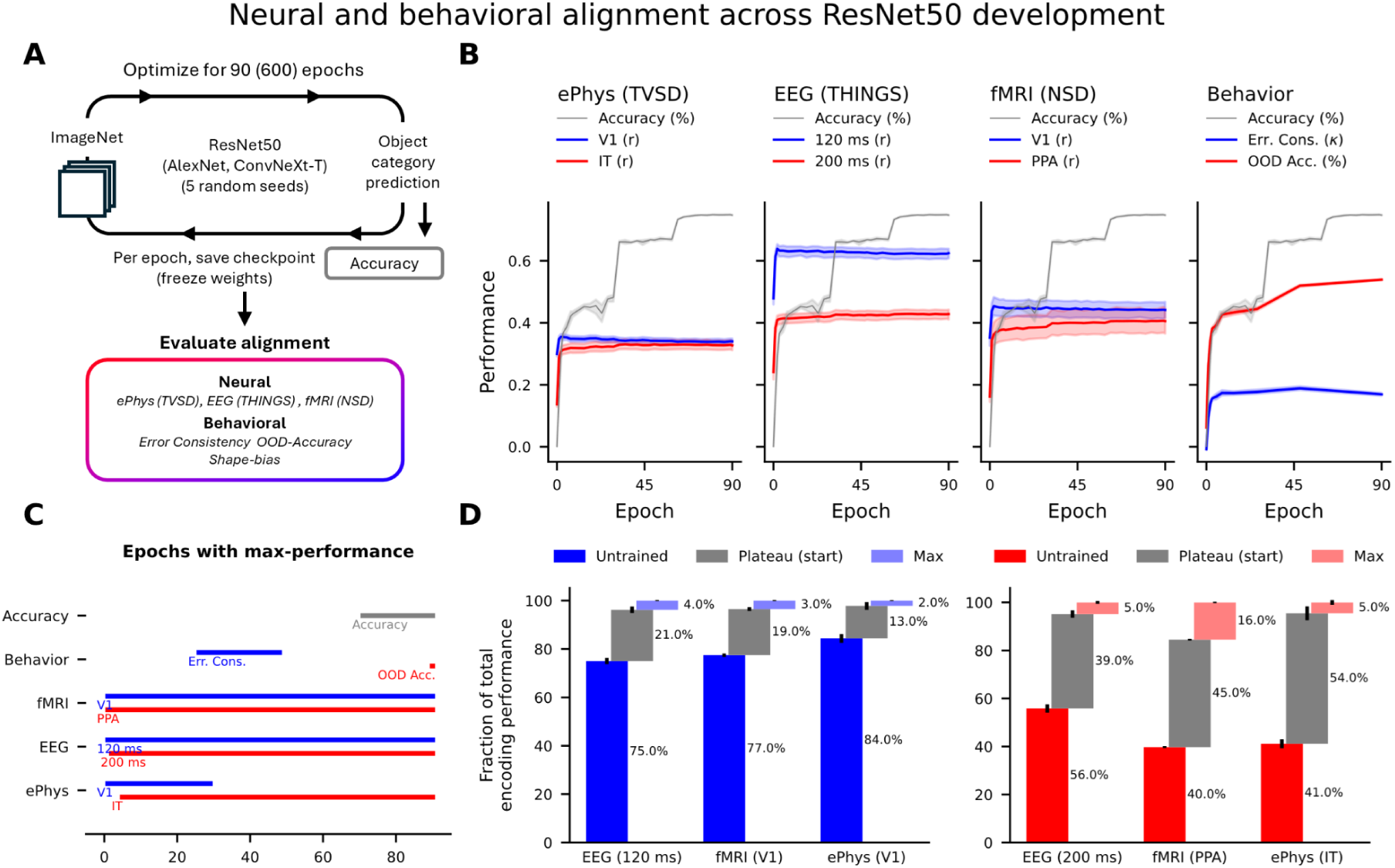
Brain alignment rises and plateaus rapidly, within the first few training epochs. **(A)** Pipeline: DCNNs trained on ImageNet from 5 random seeds (90 epochs for AlexNet and ResNet-50, 600 epochs for ConvNeXt-T); per-epoch checkpoints evaluated for classification accuracy, neural encoding across three datasets (ePhys/TVSD, EEG/THINGS, fMRI/NSD), and behavioral measures. Panels B–D show results for ResNet-50; AlexNet and ConvNeXt-T are reported in Supplementary Figure 2. **(B)** Network behavior and alignment with neural data across the training trajectory. Blue: early targets (V1 ePhys, V1 fMRI, 120 ms EEG, error consistency); red: late targets (IT ePhys, PPA fMRI, 200 ms EEG, OOD accuracy); grey: validation accuracy. Each performance measure is shown on its native 0–1 scale (encoding performance r, error consistency κ, accuracy %) and values are not comparable across measures. Shading: standard error of the mean (SEM) across subjects (encoding performance) or models (model behavior). **(C)** Epochs of maximum performance, with bars marking statistically indistinguishable epochs. **(D)** Maximum encoding performance for each of the six targets, decomposed into the fraction present in the untrained network (*Untrained*), the fraction added by the onset of the plateau (*Plateau (start)*), and the remainder gained at the epoch of maximum alignment (*Max*). Early targets on the left hand side (120 ms EEG, V1 fMRI, V1 ePhys) and late targets on the right hand side (200 ms EEG, PPA fMRI, IT ePhys). Error bars show the SEM across subjects.

A first main observation is that encoding model performance rises rapidly within a few epochs of training and subsequently plateaus, while object classification performance continues to increase throughout training (see **Figure 1B**). We quantify the epoch at which the encoding performance (i.e. its mean ± standard error across subjects) becomes statistically indistinguishable from the epoch with maximum encoding performance. This plateau is reached at epoch 1 for fMRI V1 (r = 0.44 ± 0.014) and PPA (r = 0.34 ± 0.019), EEG responses at 120 ms (r = 0.61 ± 0.008), and macaque V1 electrophysiology (r = 0.35 ± 0.005). EEG at 200 ms (r = 0.41 ± 0.008) entered this plateau at epoch 2 and macaque IT electrophysiology (r = 0.32 ± 0.014) reached it last, at epoch 5 (**Figure 1C**). Next, we quantify how much of the total brain alignment is added before and after reaching this plateau.

The size of these gains is made explicit by decomposing the maximum encoding performance per neural target into the fraction already present at initialization (epoch 0), the fraction added by the onset of the plateau, and the remainder gained at the epoch of maximum encoding performance (**Figure 1D**). For the early targets (120 ms EEG, V1 fMRI, V1 ePhys; see **Figure 1D**, left), untrained networks already reach on average 78.7% of their maximum encoding performance, ranging from 75% for 120 ms EEG to 84% for macaque V1, while the networks classify at chance (0.1%). The rise to the plateau adds another 17.6%, and less than 5% separates the onset of the plateau from the maximum. For the late targets (200 ms EEG, PPA fMRI, IT ePhys; see **Figure 1D**, right), the untrained fraction is smaller, 45.6% on average, and training contributes relatively more: 46% by the onset of the plateau and 8.7% till the maximum is reached (up to 16% for PPA). Across all measures the vast majority of DCNN-brain alignment is therefore either present at initialization or acquired within the first epochs: even macaque IT reaches 81% of its maximum encoding performance by epoch 1, when top-1 accuracy is 15.6%.

As another measure of human-network alignment next to neural predictivity, we used behavioral scores to measure the generalization abilities of DCNNs throughout training: error consistency with human observers and out-of-distribution (OOD) accuracy, which have been proposed to be more suitable for assessing human-like behavior (Geirhos et al., 2018, 2020). Error consistency follows the same pattern as the brain alignment: it reaches a plateau early and changes little thereafter, with only a nominal peak around epoch 48 (κ = 0.19 ± 0.003). In contrast, OOD accuracy kept increasing throughout training, with its peak at the final epoch.

Although the gains that follow the initial rise are small relative to between subject variance, their direction is systematic. In ResNet-50, accuracy correlated positively with encoding performance for late targets (PPA fMRI r = 0.98, macaque IT r = 0.81, 200 ms EEG r = 0.92) and negatively for early targets (V1 fMRI r = −0.56, macaque V1 r = −0.82, 120 ms EEG r = −0.48; all p < 0.001; **Supplementary Figure 3A**). In AlexNet and ConvNeXt-T all evaluated areas and samples correlated positively with accuracy (**Supplementary Figure 3B,C**). These correlations are strong because the trends are consistent over training, even though the underlying changes in overall alignment are in fact quite minimal.

### Network features stabilize early while texture-based classification continues to develop

What internal changes in the network accompany the divergence between task performance and brain alignment? To answer this, we first tracked how the network’s internal features evolve, measuring feature dissimilarity across successive training epochs (**Figure 2A**). Features change substantially between epoch 0 and 1, with a dissimilarity of r = 0.91 ± 0.01 for the images used in the EEG experiment, r = 0.95 ± 0.01 for the images used in the ePhys experiment, and r = 0.85 ± 0.02 for the images used in the fMRI experiment. Similar to brain alignment, this measure stabilizes after only a few epochs of training. However, this feature dissimilarity metric only measures the overall magnitude of feature change, but not what kind of features the network develops.

**Figure 2.**
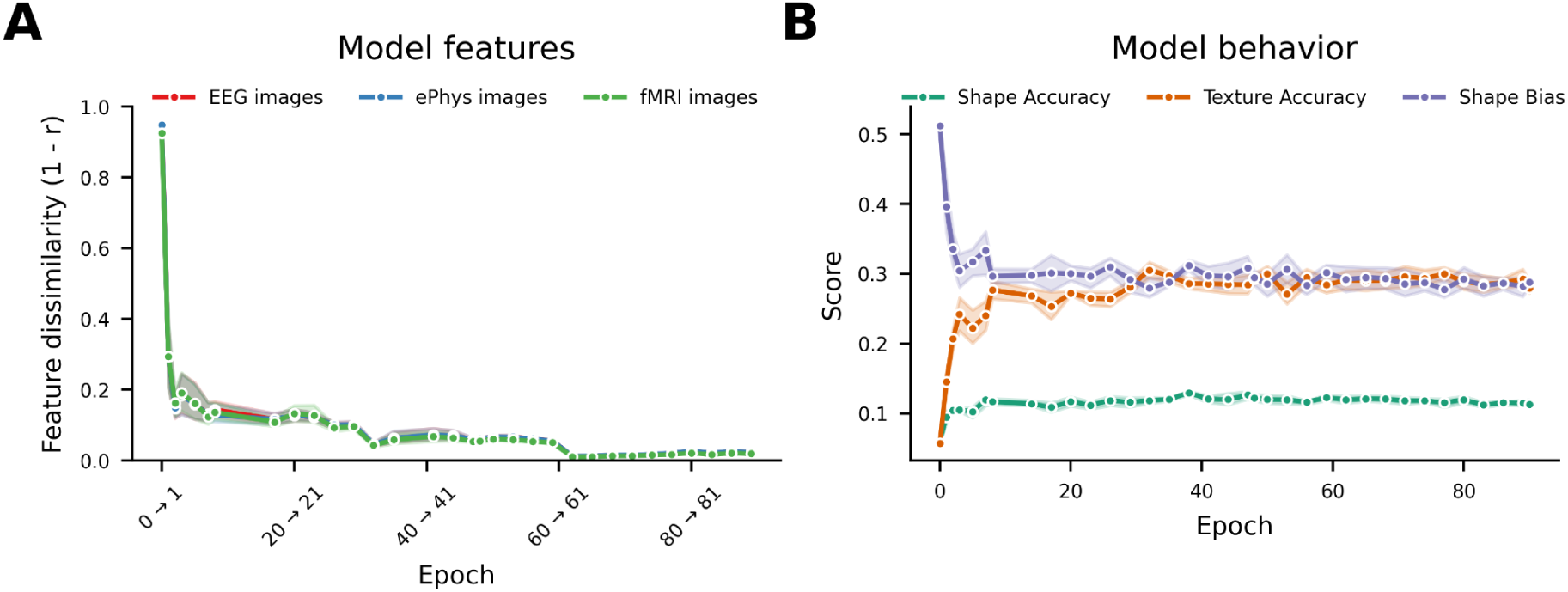
Network features stabilize early while texture-based classification continues to develop. **(A)** Feature dissimilarity (1 − r) between successive epoch pairs, computed separately for the image sets used in the EEG, ePhys, and fMRI encoding analyses. Dissimilarity is largest across the earliest transitions and small thereafter. **(B)** Shape accuracy (green), texture accuracy (orange), and shape bias (purple) on the cue-conflict test set (Geirhos et al., 2019). Shape accuracy enters its plateau by epoch 7 and texture accuracy by epoch 32; shape bias declines from 0.38 at epoch 1 to 0.27 at the end of training. Both panels: ResNet-50, n = 5 seeds; lines show the mean and shaded areas show the SEM across seeds. See Supplementary Figure 4 for results for AlexNet and ConvNeXt-T.

To characterize the kind of features underlying this early change and the subsequent stabilization, we examined the network’s classification strategy using cue-conflict stimuli that pit object shape against texture cues (Geirhos et al., 2019). Shape accuracy rises rapidly during the first few epochs of training and enters a plateau after approximately 7 epochs (**Figure 2B**). Texture accuracy develops more slowly, entering a plateau only at epoch 32. Because shape bias is the ratio of correct shape to correct shape-and-texture responses, this increase in texture accuracy naturally lowers shape bias across the first third of training after which it stabilizes. So, networks are initially more shape-biased but then adopt a texture-based classification strategy that is fully established in the first half of training, well before classification accuracy approaches its maximum. Human observers show a markedly different pattern. When presented with the same cue-conflict stimuli, humans classify by shape approximately 80 to 95% of the time (Geirhos et al., 2019). The networks never approach this value, and when fully trained have a lower shape bias compared to early checkpoints.

In sum, developmental tracking shows that encoding performance and shape accuracy enter their plateaus within the first ten epochs of training, texture accuracy only does so at epoch 32, and classification accuracy never plateaus. Brain alignment therefore develops on the same timescale as shape-based classification and on a different one from texture-based classification. Whether this shared timing reflects shared features is a separate question, which the next analyses address directly.

### Features supporting brain alignment are largely independent of features supporting task performance

The preceding analyses demonstrate that brain alignment and task performance follow different trajectories during training, with texture-based classification emerging as accuracy continues to rise. But do brain-aligned and task-relevant representations compete for the same features, or do they constitute independent systems within the network? To address this directly, we developed a lesioning analysis that quantifies the contribution of each network kernel to two measures: brain alignment and task performance.

To measure a single kernel’s contribution to each function, one might lesion the kernel in isolation and record the change in performance. But DCNN kernels do not act in isolation: their contributions depend on which other kernels are active, and single-kernel lesioning cannot capture these interactions. We therefore estimated each kernel’s contribution by lesioning random combinations. In each of 500 iterations, we set the activations of a random 50% of kernels to zero and read out both functions from the resulting activation pattern (**Figure 3A**). Because the lesioned set is drawn anew each iteration, every kernel is intact in roughly half the iterations and lesioned in the rest. Each iteration’s two scores are then assigned to the kernels that were intact in it, and averaging over iterations yields an average contribution to each function per kernel. Crucially, both readouts are taken from the same lesioned network in each iteration, so the two contribution scores are measured under identical conditions and can be compared directly. The same results are obtained for lesioning other fractions of kernels (**Supplementary Figure 6**). If task performance and brain alignment drew on the same kernels, a kernel’s contribution to one function would predict its contribution to the other, producing a positive across-kernel correlation between the two. A correlation near zero would instead indicate that the two functions rest on contributions that are unrelated across kernels.

**Figure 3.**
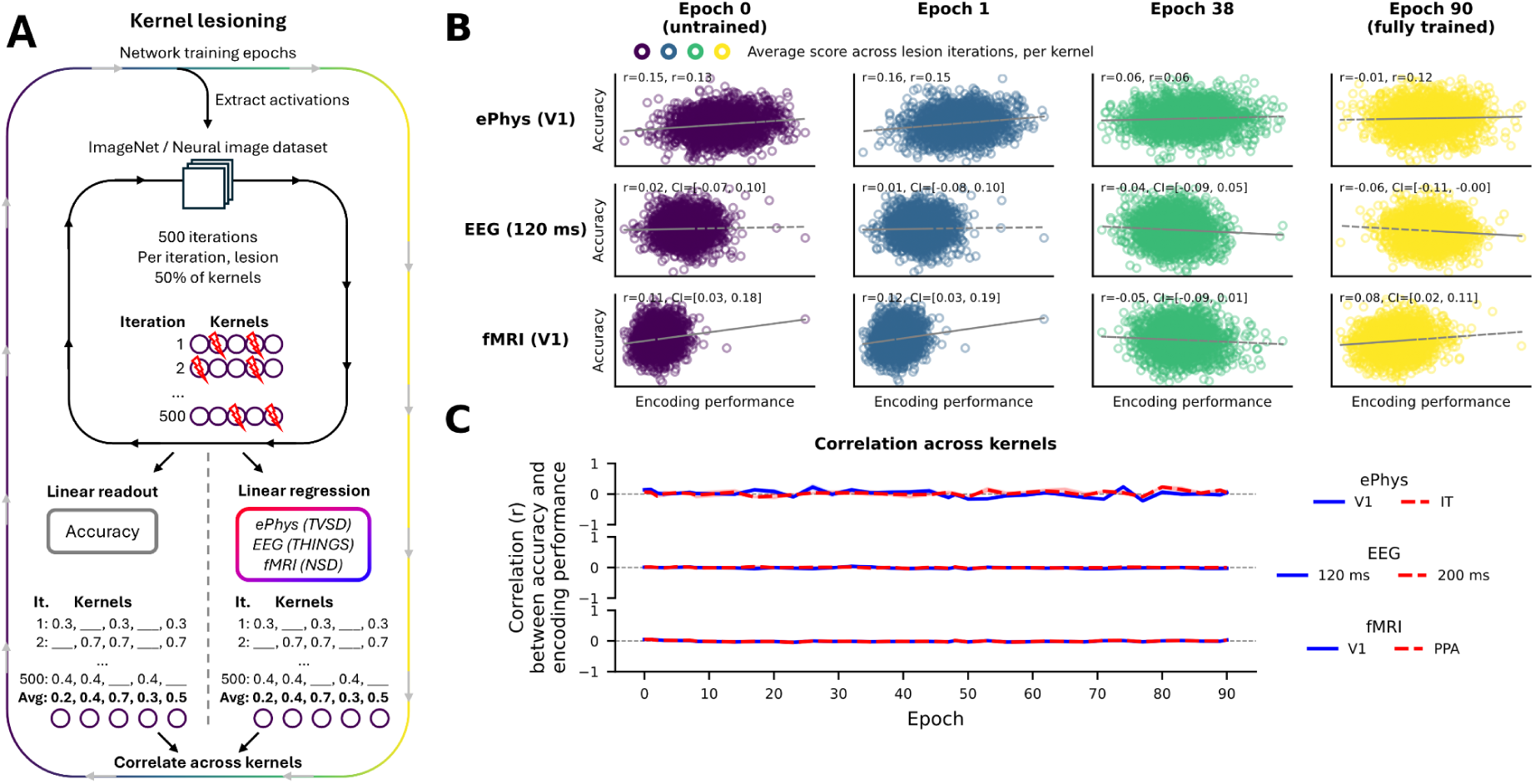
Features supporting brain encoding are largely independent of features supporting classification. **(A)** In each of 500 iterations, a random 50% of kernels are lesioned and both classification accuracy and neural encoding performance are read out from the remaining kernels; each iteration’s scores are attributed to its active kernels and averaged across iterations, yielding a per-kernel encoding contribution and accuracy contribution. These two contribution quantities are then correlated across kernels. **(B)** Encoding contribution versus accuracy contribution at representative checkpoints, shown for ePhys (V1), EEG (120 ms), and fMRI (V1); each point is one kernel. Gray dashed lines show the best linear fit and annotations show Spearman’s correlation between the contribution quantities, averaged across subjects, with 95%-confidence intervals; for the ePhys dataset the two per-monkey correlations are shown. **(C)** Spearman’s correlation between encoding and accuracy contributions across training shown separately for ePhys (V1, IT), EEG (120 ms, 200 ms), and fMRI (V1, PPA); for one example seed and averaged across subjects. Correlations remain near zero throughout training for all neural targets.

We find that the correlation is near zero at every stage of training (**Figure 3B**). At the four representative checkpoints the across-kernel correlation between the two contributions ranges from −0.05 to 0.12 for fMRI (V1), from −0.06 to 0.02 for EEG (120 ms), and from −0.01 to 0.16 across the two monkeys for ePhys (V1); none of the EEG and fMRI correlations remains significant after FDR correction, and correlations fluctuate around zero across all epochs (**Figure 3C**). A kernel’s contribution to brain alignment therefore carries essentially no, or at best a little, information about its contribution to task performance **(Figure 3B)**.

This independence holds across the visual hierarchy and across all three neural datasets (**Figure 3C**). The lack of relation is present before any training and persists across all 90 epochs, while brain alignment rises steeply in the first few epochs and task performance keeps climbing well beyond them. Even for IT, the region most associated with object recognition, this correlation remains near zero. A positive control shows that this near-zero correlation does not reflect insensitivity of the estimator: computed from the same lesioned subsets, contributions to two encoding targets within a modality (e.g., fMRI V1 and IT) are positively correlated at initialization and grade with expected feature overlap (fMRI r = 0.90, ePhys r = 0.46, EEG r = 0.10), from the substantial overlap expected between fMRI regions within a hierarchy to the limited overlap expected between EEG time windows that index distinct processing stages whereas contributions to encoding and to accuracy are not (r = 0.02 to 0.14; **Supplementary Figure 7**). Taken together, these analyses show that across kernels the contributions to brain alignment and to task performance are at most weakly correlated.

## Discussion

The alignment between deep convolutional neural networks and biological vision has been interpreted as evidence that task optimization yields brain-like computation (Yamins & DiCarlo, 2016; Schrimpf et al., 2020). Our findings challenge this interpretation. By tracking network development across the full training trajectory and comparing checkpoints with adult human and macaque vision, we observe that much of the alignment is present at initialization, and that most of what remains is established in the first few epochs. At this stage, classification accuracy is still far from its final value. After this early window, alignment no longer tracks task performance: networks continue improving at classification by developing texture-based features, but brain alignment improves at best modestly. Our results thus indicate that brain-network alignment is unlikely to reflect a shared computational solution to object recognition. Instead, we suggest that it reflects the degree to which network representations capture regularities in natural images.

Alignment is substantially in place at initialization, before the network can classify accurately at all, which suggests it reflects something more basic than a shared representation of object categories, namely features that parameterize the manifold on which natural images lie (Wright & Ma, 2022). Natural images occupy only a small region of pixel space (Pope et al., 2021) and we suggest that the coordinates of that region are not categories but the regularities that make an image natural in the first place: such images are dominated by low spatial frequencies and local correlations (Simoncelli & Olshausen, 2001), which any system operating on them would be expected to encode whether it serves recognition, navigation or action. Human vision represents this structure from the earliest stages of processing (Greene & Oliva, 2009; Scholte et al., 2009). Consistent with this, DCNNs predict human EEG responses at least as well for images that preserve local image statistics while disrupting object form as for the natural images themselves, and this alignment is decoupled from object decoding (Loke et al., in press). The regularities may be universal to natural images or they may be specific to a human sample of the world, since our ImageNet images are photographs of human environments taken by human photographers. Both locate the source of alignment in the input rather than in the objective (Bosch et al., 2026). Comparisons across fully trained networks lead to the same conclusion: the dimensions predicting the human brain best are those that recur across architectures and training objectives, and reducing a network to a small set of these leaves its alignment largely intact (Chen & Bonner, 2025). Since these networks share little beyond their training diet of natural images, the dimensions that recur across them are likely those the images impose: the coordinates of the natural image manifold. Our results suggest that these dimensions are already present within the first few training epochs, while classification accuracy is still low.

This image-structure account is consistent with our finding that alignment is established early for V1, IT and PPA alike. If alignment were carried by a category code that the brain and DCNNs share, then we would expect it to concentrate in category-selective regions and to emerge gradually with network classification performance. Neither holds in our results. We note that two of the three regions are tuned to properties of the image rather than to category: V1 neurons act as bandpass spatial frequency filters (De Valois et al., 1982), and the PPA responds to the spatial layout of a scene rather than to the objects in it (Epstein & Kanwisher, 1998). However, IT is the exception: its response patterns cluster by animacy in human and macaque alike (Kriegeskorte et al., 2008), so this is where a shared category code should show. Instead, encoding performance for IT is at 81% of its maximum by epoch 1, when classification accuracy stands at only 15.6% of an eventual 75%. That such organization can arise without recognition has been shown directly: unrecognizable texforms, which preserve object texture and coarse form, elicit the animacy topography across the ventral stream (Long et al., 2018). Even in IT, then, alignment is substantially carried by image structure that is available before, and without, object recognition by the network.

The alignment already present at initialization (**Figure 1B, D**) points to the architecture itself. The structure of a convolutional network captures a great deal of low-level image statistics before any learning (Ulyanov et al., 2018), and untrained convolutional networks predict responses in macaque and human visual cortex better than untrained transformers or fully connected networks (Kazemian et al., 2025). This advantage rests on two design choices: local connectivity and weight sharing. Both pay off only because natural images have a corresponding structure: informative relations are spatially local, and the same relations recur across the whole image. Early visual cortex is organized around the same two regularities, with local receptive fields tiled across the visual field (Hubel & Wiesel, 1962). Architecture and learning are therefore sources of brain alignment for the same reason: both depend on the structure of the input rather than the demands of the objective function. On this account, supervised networks and networks trained with an objective that never sees labels, such as reconstructing masked image patches (He et al., 2022), should acquire alignment at a similar rate over the first epochs, diverging only once training begins rewarding the features that separate categories. Alignment is known to saturate under self-supervised training as well (Gokce & Schrimpf, 2025); whether the early-epoch trajectory matches is a direct test we leave to future work.

If the early rise of brain alignment across network training indeed reflects shared natural image structure, the question is what the network learns afterwards, and why that later learning brings it no closer to human vision in either alignment or error consistency. We note that the small but consistent post-plateau gain in encoding performance, most pronounced for PPA fMRI (**Figure 1D, Supplementary Figure 2**), need not reflect increased alignment per se: linear encoding models profit from a more expressive feature basis, so predictions can improve simply because training enriches the features, making the measured gain an upper bound on the growth of alignment proper. We suggest the answer is task specialization: once the coarse structure of natural images is captured, further optimization is devoted to features that separate ImageNet categories, and these are largely not the features human vision relies on. Consistent with this, a kernel’s contribution to classification carries essentially no information about its contribution to brain alignment (**Figure 3**). The trained network’s classification rests heavily on texture (Geirhos et al., 2019), a reliance that is present from the first interpretable checkpoint and deepens over training (shape bias 0.38 at epoch 1, 0.27 at convergence), so the accuracy the network gains is obtained by refining features human vision does not weight. Human vision likely weights shape not because it solves classification better but because it was never solving only that problem: constraints such as viewpoint invariance favor shape over local texture (DiCarlo & Cox, 2007), and training on viewpoint-diverse renderings indeed moves networks toward shape-based classification (Luo et al., 2026). We speculate that this is also why error consistency does not improve: the network’s errors are driven by texture and the humans’ by shape, so refining texture features cannot bring the two sets of errors into closer agreement.

Task specialization has two further observable consequences in our data. First, in ResNet-50, the architecture with the highest average encoding performance of the three we trained, alignment with early visual targets drifts downward over training while accuracy climbs, whereas higher-area alignment stays within the plateau. The low-frequency structure that drives early alignment is acquired first because it carries the most variance (Saxe et al., 2013, 2019; Field, 1987), not because it is a training goal, and nothing in the classification objective preserves it: plausibly, as kernels are re-tuned toward features that separate categories, the ones supporting V1 alignment are overwritten. This decline is small relative to subject variance and among the three architectures we see it only here, but training beyond the first few epochs improves the model of early visual cortex in none of them: across all three, the best model of V1 is an early-epoch ResNet-50, not any fully trained network. Second, the same logic bears on scaling. Gokce and Schrimpf (2025) trained over 600 networks and found that behavioral alignment keeps improving with scale while brain alignment saturates; we vary training time with network and dataset size held fixed and find the same asymmetry. We suggest the near-zero coupling between kernel contributions may be one reason: to the extent that the features supporting alignment are separate from those supporting classification, optimizing for the task cannot be expected to improve alignment. Improving alignment may therefore require intervening on how networks learn rather than on scale alone.

Bowers et al. (2023) raise a problem for the interpretation of comparisons between networks and brains: a benchmark score, however high, leaves untested which features produce it, so good predictions may be mediated by systems that share little with biological vision. For work that treats networks as implementation models of the ventral stream, in which alignment is taken to certify a shared mechanism, this objection is serious. We suggest our lesioning analysis is one way to ask directly which features carry the prediction (**Figure 3**). What makes the question answerable is the property that separates a network from a brain: it is artificial, so any subset of units can be removed and the consequences for any readout measured directly. This points to a use of DCNNs closer to a model organism than to an implementation model (Scholte, 2018). A model organism is informative because it shares biology with us through common descent; a network shares no ancestry with the brain, and on our account its sensitivity to natural image structure is acquired independently, from the same visual world. What the network offers instead is experimental tractability: every unit is accessible and every manipulation is possible. On this view, a network is an experimental system rather than a claim about how the brain works. Training stage then joins architecture, training data, and learning rule (Yamins & DiCarlo, 2016; Conwell et al., 2024) as a variable to manipulate. By the onset of the plateau, networks have reached on average 94% of their maximum encoding performance (**Figure 1**), so a minimally trained network predicts neural responses about as well as a fully trained one. The choice of checkpoint is therefore as much a part of model selection as the choice of architecture.

Several limitations warrant acknowledgment. We study supervised classification on ImageNet, a dataset whose object-centric, non-ecological image statistics have been criticized as a poor match to human visual experience (Mehrer et al., 2021), and whose biases have been linked to the growing divergence between model accuracy and biological alignment (Linsley et al., 2026). This dependence on the training distribution is not incidental but built in: if early alignment reflects a set of images that overlaps with human visual experience by construction, our results speak only to the human-centric case, and how much alignment would survive training on images sampled outside that overlap is open (Bosch et al., 2026). The training objective is a second dependence. Under supervised classification the dissociation we report has a clear explanation: the texture features that raise ImageNet accuracy are those that diverge from biological vision. Other training objectives need not behave this way. Two observations temper this concern. Once trained, varied architectures and objectives predict brain responses comparably (Conwell et al., 2024), and Gokce and Schrimpf (2025) report that saturation persists under self-supervised training (SimCLR, DINO) and multiple architectures, though contrastive objectives would not settle the question, since their augmentations discard part of the low-level structure at issue. Our prediction that supervised and label-free networks acquire alignment at a similar rate over the first epochs remains untested. Supplementary analyses show that the core findings, early emergence, subsequent dissociation from task performance, and independent feature sets, generalize across AlexNet and ConvNeXt-T, though the trajectory for early visual areas varies between decline and plateau.

Finally, one of our major claims rests on a null result. The lesioning correlations are near zero rather than zero, so the analysis places a bound on the overlap between the two feature sets rather than excluding it. The estimator itself is sensitive enough to detect shared contributions when they exist (**Supplementary Figure 7**), so the near-zero correlation reflects the data rather than the method. That bound is informative in one direction only: overlap would not by itself have established shared computations, whereas its near-absence makes shared computations unlikely. The kernel lesioning approach also estimates average linear contributions and cannot characterize specific kernel interactions; since brain-network alignment is itself standardly measured through linear encoding models, this limitation applies to the field’s primary metric as well.

In sum, our results show that brain alignment in DCNNs is substantially in place before training begins, is nearly complete within the first few epochs, and rests on features largely independent of those supporting classification. High alignment therefore indicates that a network captures the structure of natural images, not that it has arrived at the brain’s solution to object classification. Improvements in classification performance will accordingly not be the full answer to improving brain alignment; that will require intervening on how networks learn. One lever is the input distribution: developmentally inspired training diets improve shape bias and robustness (Vogelsang et al., 2025; Lu et al., 2026; Müller, Snoek, et al., 2026), though whether they also improve alignment is unknown. Another is the objective itself: our lesioning analysis showed that the features carrying encoding contribution receive no reward from classification, so an objective that optimizes neural prediction (Seeliger et al., 2021; El-Gazzar & Van Gerven, 2025) would reward the features classification leaves untouched.

## Methods

We evaluated representational alignment between deep convolutional neural networks (DCNNs) and human and macaque neural responses as well as behavioral data across DCNN training, for three neural datasets, three network architectures, and five network seeds.

### Network Training and Architecture

Three architectures, ResNet-50 (He et al., 2016), AlexNet (Krizhevsky et al., 2012), and ConvNeXt-T (torchvision v1 training recipe; Liu et al., 2022) were trained on ImageNet-1k object classification (ILSVRC 2012) for 90 epochs (ResNet-50, AlexNet) or 600 epochs (ConvNeXt-T) using stochastic gradient descent with momentum = 0.9, weight decay = 1 × 10⁻⁴, batch size = 256, and cross-entropy loss. The initial learning rate was 0.1, reduced by a factor of 0.1 every 30 epochs. Images were first resized to 256 × 256 pixels and then a central crop of 224 × 224 pixels was taken. Training was performed in PyTorch 2.2.0 on NVIDIA A100 GPUs. Network checkpoints were saved at multiple epochs (dense sampling for early epochs, sparse sampling for later epochs) to track changes throughout training. Each of the three architectures was instantiated with five random seeds to account for variability due to initialization. Results for ResNet-50 are presented in the main figures; AlexNet and ConvNeXt-T results are presented in **Supplementary Figures 1–5**.

For each checkpoint, we extracted activations from the final layer of each residual block (blocks 1–4) for ResNet-50, from all ReLU layers (features.1, features.4, features.7, features.9, features.11) for AlexNet, and from the final convolutional layer of each block (features.0, features.2.1, features.4.1, features.6.1) for ConvNeXt-T, providing a hierarchical sampling of network representations. Activations were extracted for all stimuli in each neural dataset.

### Neural Datasets

#### Macaque Electrophysiology

We used the THINGS Ventral-stream Spiking Dataset (TVSD; Papale et al., 2025), which contains multi-unit activity recordings from two macaque monkeys across visual areas V1, V4, and IT. Recordings were obtained using chronically implanted multi-electrode arrays while the animals performed a passive fixation task. The stimulus set comprised over 25,000 natural images from the THINGS database (Hebart et al., 2019), presented for 100 ms each with 100 ms inter-stimulus intervals. Neural responses were quantified as spike counts in a 70–170 ms window post-stimulus onset. For the encoding analysis, we used the individual time-resolved responses of electrodes in V1 and IT.

#### Human EEG

We used the large-scale EEG dataset from Gifford et al. (2022), comprising recordings from 10 participants who each completed approximately 82,160 trials across 16,740 unique image conditions from the THINGS database. EEG was recorded using a 64-channel system at 1000 Hz, downsampled to 100 Hz for analysis. We focused on two time windows capturing distinct processing stages: early responses (120 ms post-stimulus), reflecting the initial feedforward sweep through visual cortex (Cichy, Pantazis, & Oliva, 2014), and late responses (200 ms), reflecting recurrent and higher-level processing.

#### Human fMRI

We used the Natural Scenes Dataset (NSD; Allen et al., 2022), a large-scale 7T fMRI dataset comprising responses from 8 participants who each viewed up to 10,000 natural scene images over 30–40 scanning sessions. We focused on the 872 "shared" images viewed by all participants, using the single-trial beta estimates preprocessed in 1.8mm volume space and denoised using GLMdenoise (betas_fithrf_GLMdenoiseRR). Betas were z-scored within each scanning session and averaged across repetitions. For the encoding analysis, we used the individual voxel responses in the NSD-defined Regions of Interest (ROIs) V1 and PPA (NSD contains no localizer for the object-specialized lateral occipital complex (LOC) which would have been the ideal neural target to compare object-trained networks with).

### Brain Alignment Quantification

#### Encoding Models

We quantified network-brain alignment using linearized encoding models that predict neural responses from network activations. For each network seed, checkpoint, and layer, we did the following: we first extracted the activations for each stimulus image for each of the three neural datasets. Per stimulus set, we then performed a Principal Component Analysis (PCA) across the features, reducing the number of components to 100. Subsequently, we fit a linear regression model to predict neural responses (per participant and voxel for fMRI, per participant, electrode, and time point for EEG, per subject, electrode, and time point for electrophysiology) from network activations.

For the EEG and electrophysiology datasets, we used the pre-defined training and test splits of stimuli and neural data to fit both the PCA and the linear regression models. For fMRI data, we used 5-fold cross-validation across images, fitting the PCA and regression model on 80% of stimuli and evaluating on the held-out 20%. We report the cross-validated encoding performance, which was quantified as the Pearson correlation between predicted and actual responses on the held-out test sets, respectively. Encoding performance scores were averaged per participant across voxels within ROIs V1 and PPA for fMRI, and per participant and time point (120 and 200 ms) across electrodes for EEG. For electrophysiology, we averaged encoding performances for each subject across electrodes in V1 across all time points between 0 and 100 ms and in IT across all time points between 100 and 200 ms. Additionally, we averaged encoding performances across network seeds and layers, for each network architecture.

Since the TVSD dataset only consists of two subjects, we used bootstrapping across neurons to quantify the reliability of the encoding performance, to calculate the overlap of the error bands in **Figure 1C**. Specifically, for each subject and each ROI (V1, IT), with N neurons per ROI, we randomly drew (with replacement) N neurons and averaged the encoding performance across those neurons and across subjects, then determined the SEM per epoch across bootstrap iterations and subsequently evaluated per epoch whether the SEM overlaps with that of the epoch with maximum encoding performance.

### Representational Similarity Analysis

For the feature similarity analysis (**Figure 2A**), we computed representational dissimilarity matrices (RDMs) for each checkpoint and layer. Each RDM captured pairwise dissimilarities (1 − Pearson correlation) between activation patterns for all images in the stimulus set (except for fMRI for which we randomly sampled 16,000 stimuli from the set of all stimuli across participants). We then computed the correlation between RDMs from different epochs to quantify representational similarity across training. Feature dissimilarity was computed as 1 − *r* for successive epoch transitions.

### Behavioral Measures

#### ImageNet Classification Performance

We evaluated top-1 classification accuracy on the ImageNet-1k validation set (50,000 images) at each training epoch. This provided the primary measure of task performance.

#### Shape-Texture Classification Dynamics

We assessed shape versus texture bias using the cue-conflict stimulus set from Geirhos et al. (2019). These stimuli are created by applying the texture of one object category (e.g., elephant skin) to the shape of another (e.g., cat silhouette), forcing a choice between shape-based and texture-based classification. For each network checkpoint, we computed three measures: (1) Shape accuracy: the proportion of cue-conflict images classified according to their shape; (2) Texture accuracy: the proportion classified according to their texture; (3) Shape bias: Shape accuracy / (Shape accuracy + Texture accuracy), ranging from 0 (pure texture bias) to 1 (pure shape bias).

#### Out-of-Distribution Accuracy

We evaluated generalization to distribution-shifted images using the stimulus sets from Geirhos et al. (2018), which include images with manipulations such as high-pass filtering, low-pass filtering, contrast reduction, and phase scrambling. OOD accuracy was computed as mean classification accuracy across all manipulation conditions.

#### Error Consistency

To assess human-model behavioral alignment, we computed error consistency (Cohen’s kappa) between models and human observers following Geirhos et al. (2020). For each image we recorded whether each system classified it correctly or incorrectly, giving a binary correct/incorrect response per system. We then computed the observed agreement, c_obs, as the fraction of images on which model and human responses coincided (both correct or both incorrect), and the agreement expected by chance from the two accuracies, c_exp = p_model · p_human + (1 − p_model)(1 − p_human). Error consistency is the chance-corrected agreement, κ = (c_obs − c_exp) / (1 − c_exp). Values above zero indicate that model and human decisions coincide on the same individual images more often than their accuracies alone predict; values near zero indicate agreement no greater than chance. Because the chance correction removes the contribution of overall accuracy, the measure reflects trial-by-trial agreement on which images are difficult rather than how many are classified correctly.

### Kernel Lesioning Analysis

To directly assess whether the same network features support brain encoding and classification, we performed lesioning experiments that quantify the average contribution of each convolutional kernel to both ImageNet accuracy and encoding performance for the three neural datasets (**Figure 3**). Relating per-kernel contributions to two functions follows Scholte et al. (2018), who measured each unit’s contribution to two tasks via marginalization of single units. Because DCNN kernels interact, we adopted a random sampling approach that allows estimation of the average contribution of individual kernels despite the combinatorial extent of interactions between kernels; similar strategies have been developed in lesion analysis (Fakhar & Hilgetag, 2022) and explainable AI (Petsiuk et al., 2018).

### Lesioning

For each network architecture, seed, and training checkpoint, we first extracted DCNN activations for 20,000 random ImageNet images (20 per category) and the stimuli from the three neural datasets. We split the ImageNet activations into training (80%) and test (20%) splits and also use the same train and test splits for the neural datasets as described earlier (see Methods section Encoding Models). Before lesioning, we first fit a linear classifier once on the intact activations from the ImageNet training set. Likewise, we fit an encoding model (PCA plus linear regression) once on the intact activations of the training sets for each neural dataset and neural target as described earlier.

Across 500 iterations, we then randomly lesioned a fixed fraction of kernels and assessed both ImageNet test accuracy and cross-validated encoding performance for each dataset. Specifically, in each iteration, we randomly selected 50% of kernels and set their extracted activations to 0 across all test stimuli (other lesion fractions in **Supplementary Figure 6**). We then used the fitted ImageNet classifier and the fitted encoding models to evaluate performances (test accuracy for ImageNet classification and cross-validated encoding performance for each neural target) on the lesioned activations of the test set. Because the readout weights are fixed while kernel activations are zeroed at evaluation, the contribution scores measure the network’s reliance on each kernel under the trained readout, rather than the information that would remain after refitting. Across iterations, we tracked accuracy and encoding performances and assigned these to the kernels that were *not* lesioned, thus accumulating the scores per kernel across many iterations, and finally averaged accuracy and encoding performance per kernel. These per-kernel scores correspond to Banzhaf values (Banzhaf, 1965).

### Contribution Analysis

We computed Spearman’s correlation between encoding contribution and accuracy contribution across kernels. A positive correlation would indicate that features supporting brain encoding also support classification; a correlation near zero would indicate that different features serve each function. We performed this analysis separately for participants, voxels, electrodes, and time points and averaged results with V1 and PPA (fMRI), V1 and IT (electrophysiology), and for 120 and 200 ms (EEG) to assess consistency across the visual hierarchy. In **Figure 3**, and **Supplementary Figures 5, 6,** we report the correlation between per-kernel encoding and accuracy contributions, computed separately for each subject; we report the average correlation across subjects together with 95% confidence intervals derived from the across-subject distribution of these correlations (n = 8 subjects for fMRI, n = 10 for EEG). For the electrophysiology dataset, with two animals, we report the correlation per monkey without a confidence interval or p-value. Additionally, we obtained p-values using a permutation test with 1,000 iterations and corrected for multiple comparisons using the Benjamini-Hochberg false discovery rate procedure. As a positive control, we correlated the contributions to the two different encoding targets within each modality (V1 and IT; 120 and 200 ms; V1 and PPA), computed from the same lesioned subsets, at initialization (**Supplementary Figure 7**).

## Data and Code Availability

All neural datasets used in this study are publicly available: the Natural Scenes Dataset (Allen et al., 2022; https://naturalscenesdataset.org), the THINGS-EEG dataset (Gifford et al., 2022), and the THINGS Ventral-stream Spiking Dataset (Papale et al., 2025). Network training code, analysis scripts, and trained model checkpoints will be made available upon publication on OSF (https://osf.io/g9qbf).

## Acknowledgements

Marcel van Gerven is supported by the Dutch Brain Interface Initiative (DBI2) with project number 024.005.022 of the research programme Gravitation which is cofinanced by the Dutch Research Council (NWO). Niklas Müller was supported by the University of Amsterdam Data Science Center Interdisciplinary PhD program. We thank Ole Jürgensen for training the five ConvNeXt-T networks, originally in the context of another project, that we use here.

## Author Contributions

H.S.S. and M.A.J.v.G. conceptualized the study. J.S. and H.S.S. performed the original analyses. J.S. trained the AlexNet and ResNet-50 networks. N.M. performed the full analyses. H.S.S. wrote the initial manuscript. N.M. created the figures. N.M., I.I.A.G., M.A.J.v.G., and H.S.S. edited the manuscript and contributed to the interpretation of the results.

---

**Supplementary Figure 1.**
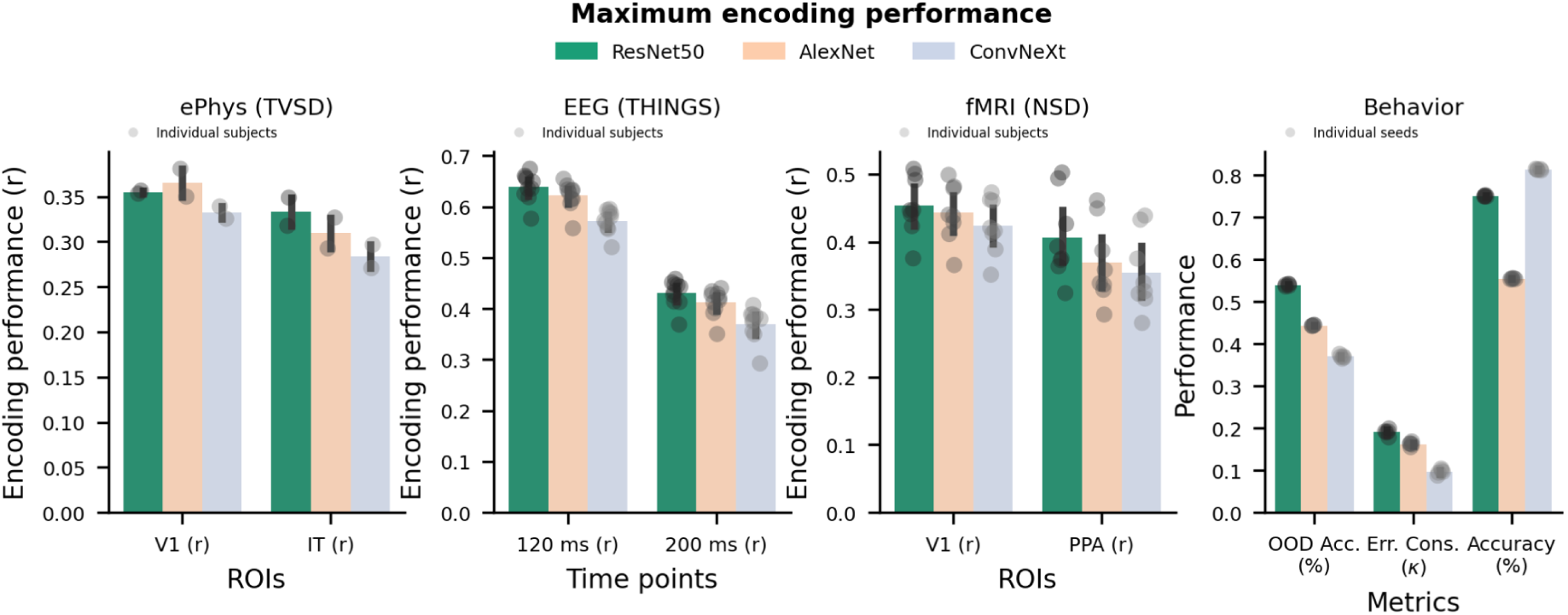
Brain-network alignment and behavioral measures are comparable across architectures. Encoding performance for the three neural datasets and two neural targets each as well as network behavioral measures, in the following order (from left to right): macaque electrophysiology for V1 and IT; human EEG at 120 ms and 200 ms; human fMRI for V1 and PPA; out-of-distribution accuracy, error consistency with human observers, and top-1 ImageNet accuracy. Colored bars show ResNet-50 (green), AlexNet (orange) and ConvNeXt-T (blue). For the encoding panels, gray dots show individual subjects, and error bars show the SEM across subjects, computed on values first averaged across the five network seeds; for the behavioral panel, gray dots show individual seeds, and error bars show the SEM across the five seeds.

**Supplementary Figure 2.**
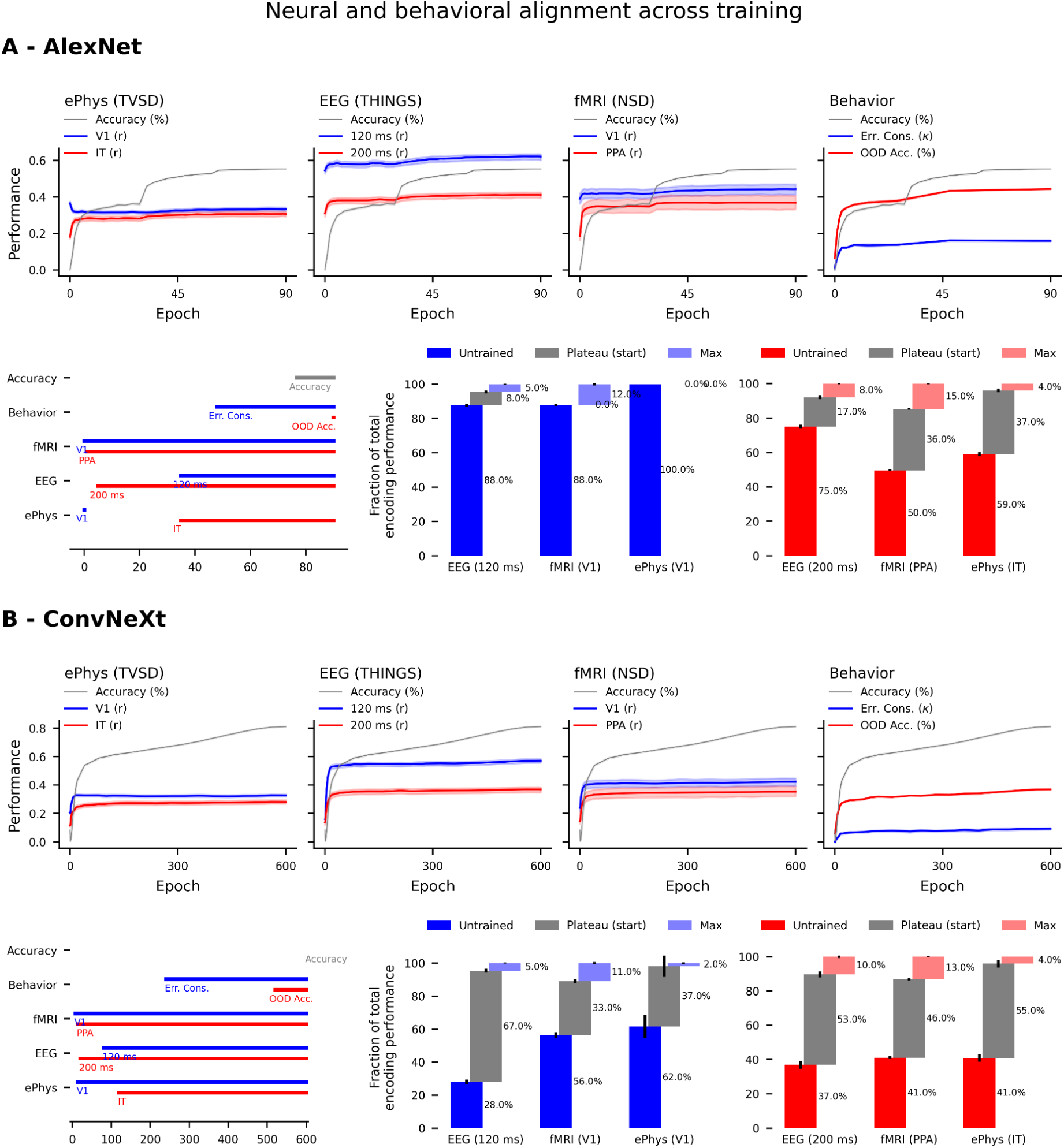
Task performance versus brain alignment for **(A)** AlexNet and **(B)** ConvNeXt-T (600 training epochs). Top panels show ImageNet validation accuracy (gray), and cross-validated encoding performance per neural dataset and target for each training checkpoint (epoch). Bars in bottom left panels indicate the epochs at which performance is indistinguishable from maximum performance across epochs. Bottom right panels show the fraction of total encoding performance present at initialization (*Untrained*), at the start of the plateau, i.e. the start of the bar in the left panel (*Plateau (start)*), and at the epoch with max performance (*Max*). For more details see Figure 1 caption and Methods. Despite lower overall encoding performance, both architectures show a similar dissociation between task performance and brain alignment.

**Supplementary Figure 3.**
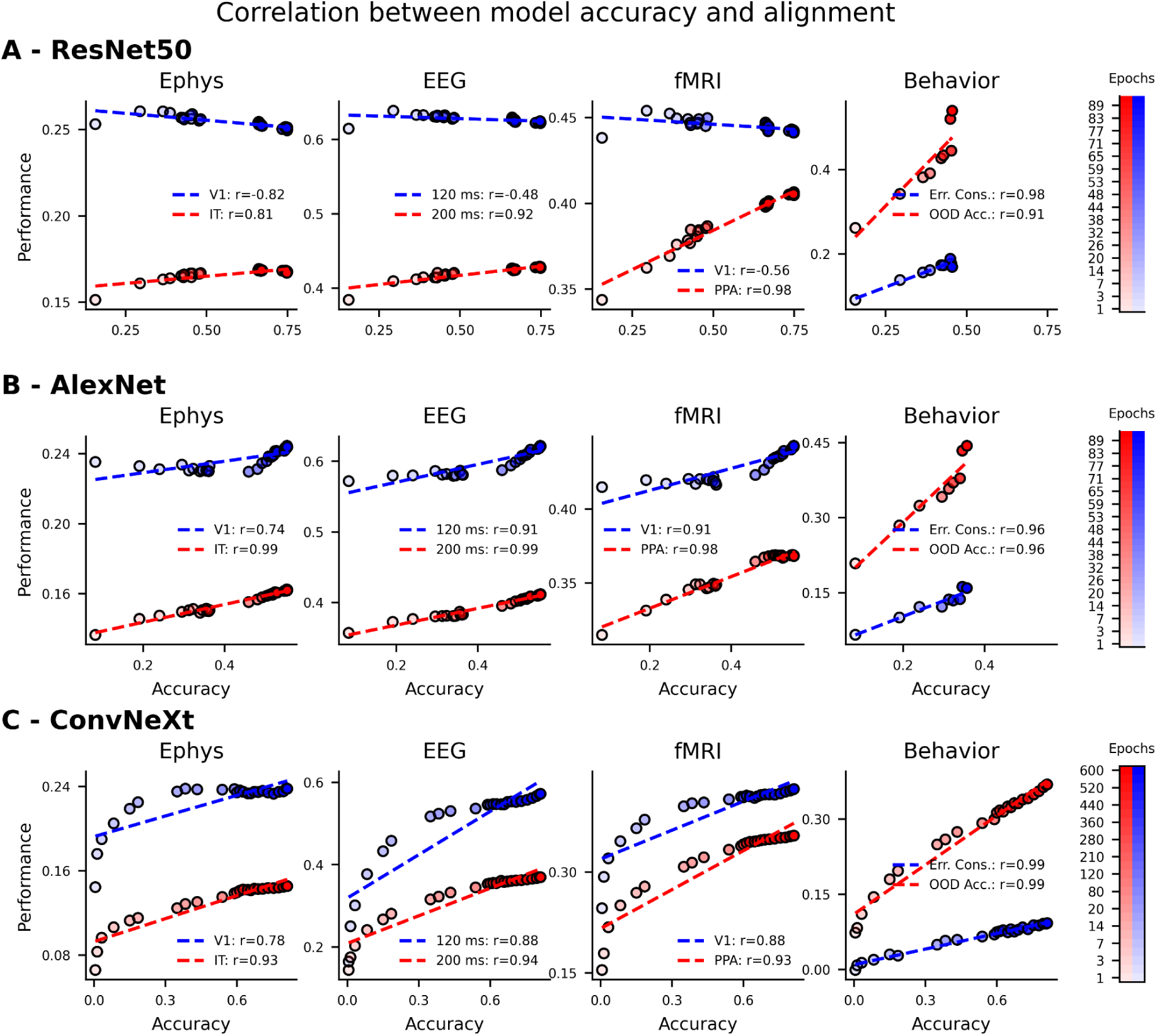
Encoding performance as a function of classification performance across training. **(A)** ResNet-50. **(B)** AlexNet. **(C)** ConvNeXt-T. Within each panel, encoding performance for macaque ePhys, human EEG and human fMRI, and the behavioral measures, are plotted against ImageNet validation accuracy across epochs, from left to right. Blue: early targets (V1, 120 ms EEG, error consistency); red: late targets (IT, PPA, 200 ms EEG, OOD accuracy). Each point is one checkpoint, averaged across five network initializations; fill runs from white at epoch 0 to full color at the final epoch. Each measure is on its native 0–1 scale and values are not comparable across measures. Dashed colored lines indicate the best linear fit and correlation values for each panel and neural target are shown in the legend. Significance of the correlations between task performance and brain alignment was assessed against the null hypothesis of no correlation using a t-test on the correlation coefficient with n − 2 degrees of freedom.

**Supplementary Figure 4.**
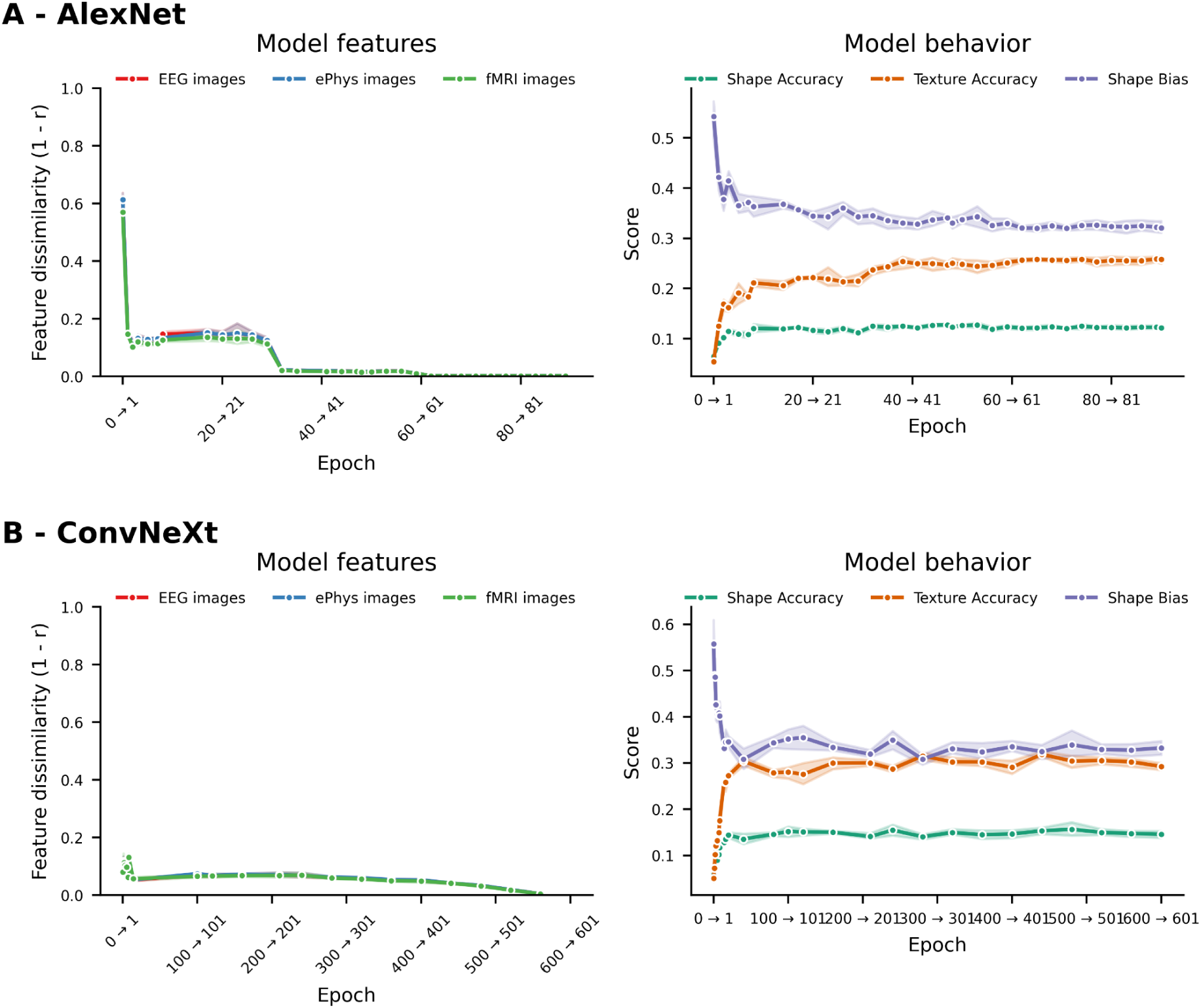
Feature stabilization and texture reliance in AlexNet and ConvNeXt-T. Panels show **(A)** AlexNet and **(B)** ConvNeXt-T, each in the format of Figure 2. Left: feature dissimilarity (1 − r) between successive epoch pairs, computed separately for the stimulus sets used in the EEG, ePhys, and fMRI encoding analyses. Right: shape accuracy (green), texture accuracy (orange), and shape bias (purple) on the cue-conflict set (Geirhos et al., 2019). n = 5 network seeds; lines show the mean and shaded areas the SEM across seeds. As in ResNet-50, features stabilize within the first few epochs while texture reliance continues to increase.

**Supplementary Figure 5.**
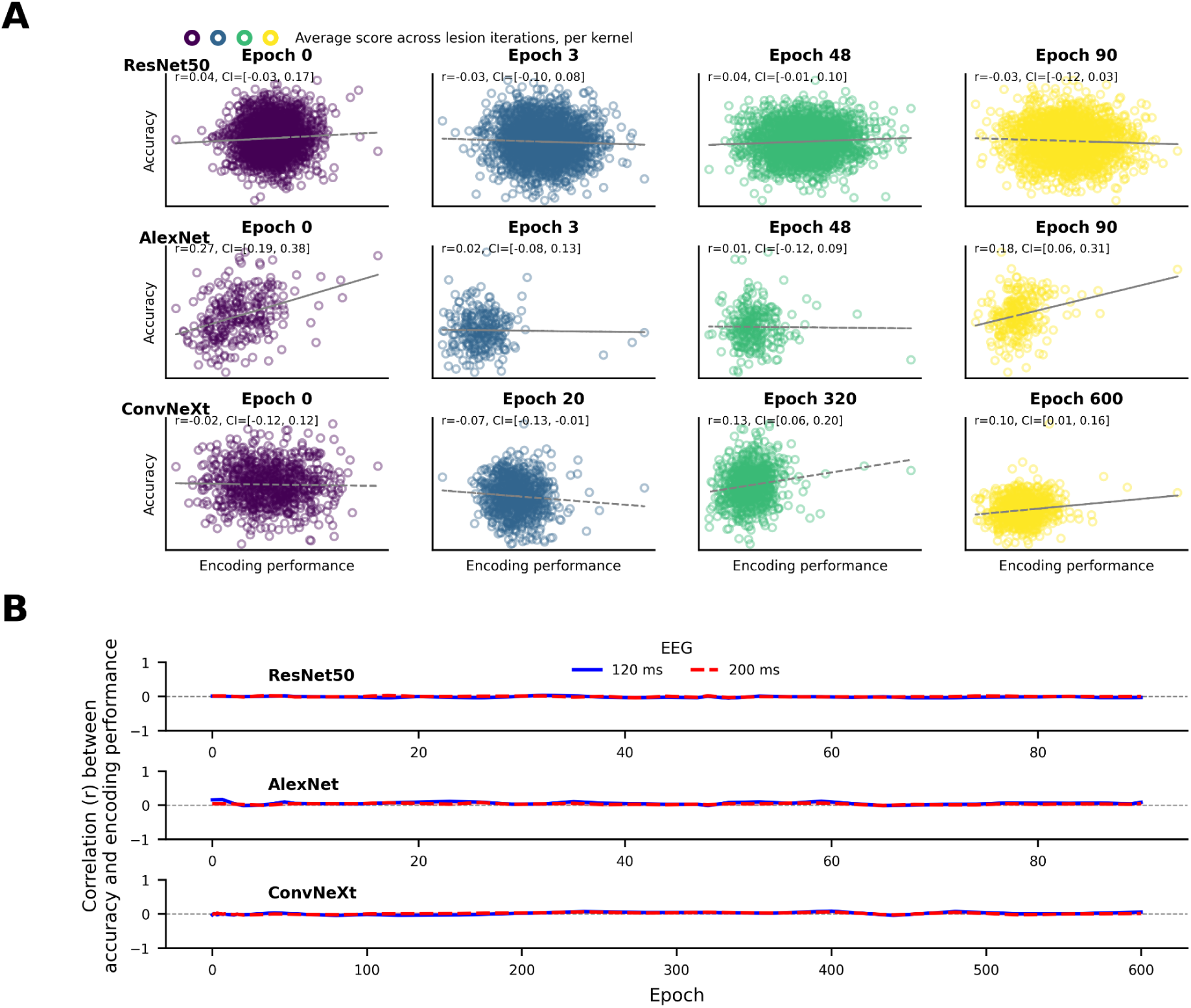
Encoding and classification contributions are independent across architectures. **(A)** Encoding contribution versus accuracy contribution for individual kernels at a lesion fraction of 0.5, for the EEG dataset (exemplary; similar results are obtained for the other dataset but not shown here), for one example seed; rows show ResNet-50 (top), AlexNet (middle) and ConvNeXt-T (bottom); columns show epochs 0, 3, 48, 90 for ResNet-50 and AlexNet, and epochs 0, 20, 320, 600 for ConvNeXt-T (all corresponding to 0%, 3%, 53% and 100% of training progress). Each point is one kernel and shows the average contribution across 500 lesion iterations, averaged over subjects, and annotations show Spearman’s correlation between the contribution quantities with 95%-confidence intervals. **(B)** Correlation between encoding and accuracy contributions across all training epochs shown separately for each architecture and each neural target for the EEG dataset. Correlations remain near zero for all three architectures throughout training.

**Supplementary Figure 6.**
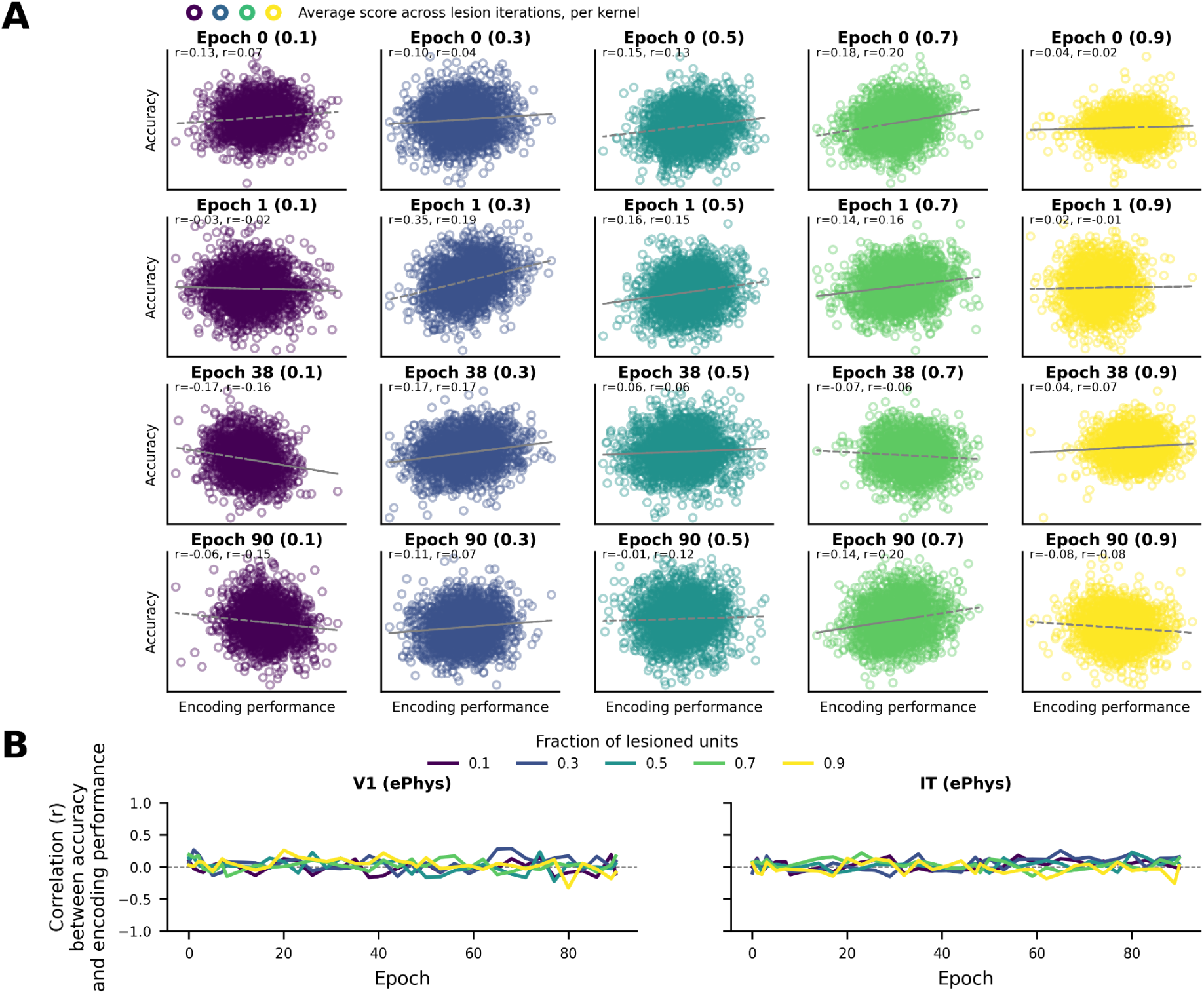
The independence of encoding and classification contributions does not depend on the lesion fraction. **(A)** Encoding contribution versus accuracy contribution for individual kernels, computed from the macaque electrophysiology data for ResNet-50, for one example network seed; columns show lesion fractions of 0.1, 0.3, 0.5, 0.7 and 0.9; rows show epochs 0, 1, 38 and 90. Each point is one kernel and shows the average contribution across 500 lesion iterations, and annotations show the two per-monkey Spearman’s correlations between the contribution quantities. **(B)** Correlation between encoding and accuracy contributions across training, from epoch 0 to epoch 90, for V1 (left) and IT (right), shown separately for each lesion fraction averaged across the two monkeys.

**Supplementary Figure 7.**
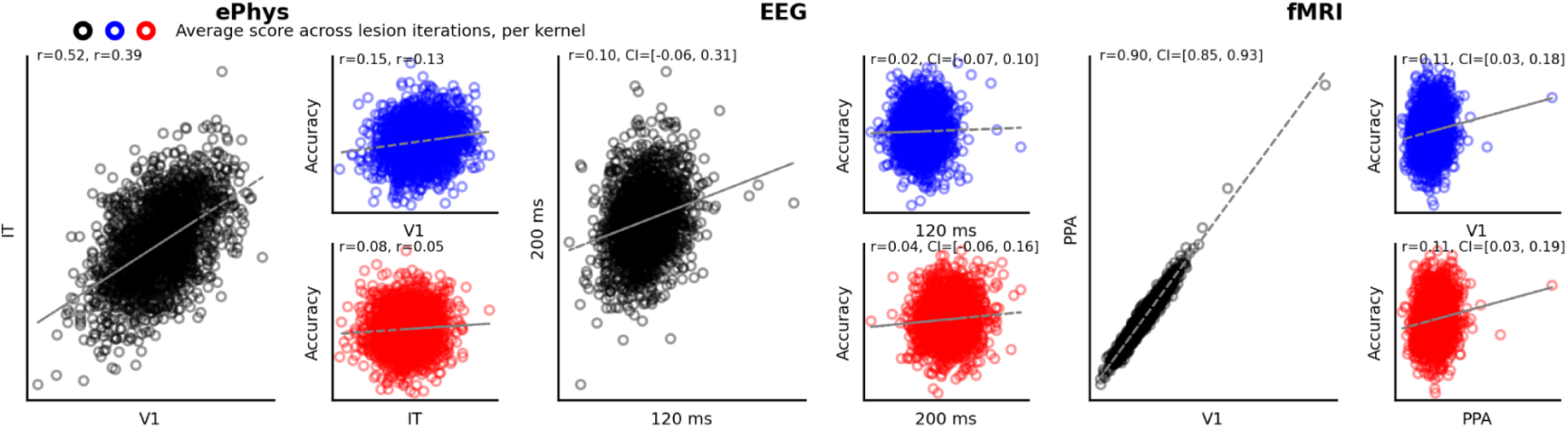
Control analyses for the kernel contribution correlations. Per-kernel contribution scores from the same lesioned subsets as Figure 3, at initialization. Each point is one kernel and shows the average contribution across 500 lesion iterations, and annotations show Spearman’s correlation (averaged over subjects) between the contribution quantities with 95%-confidence intervals; for the ePhys dataset the two per-monkey correlations are shown. Black: contributions to two encoding targets within a modality (ePhys: V1 and IT; EEG: 120 and 200 ms; fMRI: V1 and PPA). Blue and red: contributions to classification accuracy against each of those encoding targets (similar to data shown in Figure 3). All comparisons within a panel use identical lesioned subsets, so differences between them cannot reflect differences in sampling. Contributions to two encoding targets within a modality are positive and grade with expected feature overlap: highest for fMRI V1 and PPA (r = 0.90), intermediate for ePhys V1 and IT (r = 0.46, averaged over monkeys), and lowest for the two EEG time points (r = 0.10), consistent with the largely distinct neural activity captured at these two latencies.

